# Screening the Medicines for Malaria Pandemic Response Box chemical library on *Caenorhabditis elegans* identifies re-profiled candidate anthelmintic drug leads

**DOI:** 10.1101/2022.08.10.503491

**Authors:** Marina Nick, Frederick A Partridge, Ruth Forman, Carole JR Bataille, Kathryn J Else, Angela J Russell, David B Sattelle

## Abstract

The 3 major classes of soil transmitted helminths (whipworm, hookworm and Ascaris) affect 1.5 billion people worldwide mostly in poor countries, where they have adverse effects on child development, nutrition, and the work capacity of adults. Although there are drugs effective on Ascaris, notably the benzimidazoles, those same drugs show poor efficacy particularly against whipworm (*Trichuris trichiura*) and to a certain extent hookworm. Parasitic nematodes also infect farm livestock and companion animals. Resistance to currently deployed human and veterinary anthelmintic drugs is a growing problem. Therefore, new chemical anthelmintic lead compounds are urgently needed. One of the fastest routes to a novel therapeutic lead is to screen libraries of drugs which are either already approved for human use or have already been part of clinical trials. We have pursued this approach to anthelmintic lead discovery using an invertebrate automated phenotyping platform (INVAPP) for screening chemicals and the well-established nematode genetic model organism *Caenorhabditis elegans*. The 400 compound Medicines for Malaria Pandemic Response Box library was screened with each compound tested initially at 1.0 × 10^−4^ M. We identified 6 compounds (MMV1593515 (vorapaxar), MMV102270 (diphyllin), MMV1581032 (ABX464), MMV1580796 (rubitecan), MMV1580505 and MMV1593531) active in both an L1-L4 growth / motility assay and in an L4 motility assay. For vorapaxar, an EC_50_ of 5.7 × 10^−7^ M was observed, a value comparable to some commercial anthelmintics. Although not a parasite, the ease with which high-throughput screens can be pursued on the free-living nematode *C. elegans* makes this a useful approach to identify chemical leads and complement the often lower-throughput experiments on parasitic nematode models.

## Introduction

The major human gastrointestinal tract parasites collectively known as soil transmitted helminths (STHs) are the whipworm (*Trichuris trichiura*), the roundworm (*Ascaris lumbricoides*), and hookworms (*Necator americanus and Ancylostoma duodenale*). Over one billion people are estimated to be infected with at least one STH (Elfawal et al., 2019). STH infection is a significant but neglected cause of morbidity (Hotez and Kamath, 2009; Else et al., 2020). According to Hotez and Kamath (2009) more than 50 million school-aged children and 7 million child-bearing-age women in Sub-Saharan Africa are infected with one or more STH (Hotez and Kamath, 2009). Due to the remoteness and inaccessibility of the worst-affected places, the worldwide burden of STHs has most likely been underestimated (Moser et al., 2019).

Currently, preventive chemotherapy, based on single-dose, mass drug administration (MDA), is used to treat STHs in endemic regions, with the goal of minimising morbidity in pre-school and school-aged children with moderate to heavy infections. Such treatments, which use the benzimidazole drugs albendazole and mebendazole remain effective against ascariasis but less so against hookworm and whipworm. (Else et al., 2020; Moser et al., 2017, 2019; Speich et al., 2015; World Health Organization, 2019). Therefore, treatment for the STHs is an example of an unmet clinical need that necessitates the development of novel therapeutics. Recent experience from the animal health field showed that resistance to the anthelmintic monepantel (Kaminsky et al., 2008) appeared within two years of its introduction (Bartley et al., 2015). Therefore, current MDA programmes may lead to similar selection for resistance, and the efficacy of, for example, imidazole drugs against *T. trichiura* has fallen in recent years (Adegnika et al., 2015; Partridge et al., 2021).

Despite the long history of anthelmintic medication development, the majority of currently used anthelmintics were discovered by phenotypic screening of chemical candidates. Medicines that act on ligand-gated ion channels (Lees et al., 2012) and drugs that target the cytoskeletal protein tubulin are among the products of this method (Sharma and Abuzar, 1983). In the search for new anthelmintic drug candidates, several laboratories have developed high-throughput screening methods (Partridge et al., 2020). In our laboratory an invertebrate automated screening platform (INVAPP) and the algorithm Paragon (Partridge et al., 2018a) have been developed. This system has been used to identify two novel chemical classes of anthelmintics, the dihydrobenzo[e][1,4]oxazepin-2(3H)-ones (Partridge et al., 2017, 2021) and the diaminothieno[3,2-d]pyrimidines (Partridge et al., 2018b). We have therefore adopted the INVAPP / Paragon system to explore a new chemical library. *C. elegans* is a nematode genetic model organism (Brenner, 1974) that is frequently employed in anthelmintic discovery research (Fraser et al., 1999; Roy et al., 2002; Kaminsky et al., 2008; Holden-Dye et al., 2013; Burns et al., 2015; Partridge et al., 2020). The Pandemic Response Box is a chemical library supplied by Medicines for Malaria Venture (MMV) and the Drugs for Neglected Diseases Initiative (DNDi). It includes 400 drug-like compounds that are either already on the market or in various stages of research for uses other than anthelmintics. The library was selected by experts from Academia and Industry with the aim of pursuing an Open Source approach to discovering new chemical leads that could impact on neglected, pandemic-scale diseases with unmet clinical needs (Samby et al., 2022). The library contains 201 antibacterial compounds, 153 antivirals, and 46 antifungals. Compounds of interest from this library may therefore offer a fast-track route into clinical trials in the search for new anthelmintics. We use INVAPP to assess the activities of all 400 drugs on *C. elegans* L1-L4 stages using a growth / motility assay and on the L4 stage utilising a motility assay in order to understand the potential of library compounds for re-purposing as candidate anthelmintic drug leads.

## Methods

### The Pandemic Response Box Library

The Pandemic Response Box library was supplied by the Medicines for Malaria Venture. All stock compounds were supplied as 10^−2^ M stocks in DMSO.

### *C. elegans* – maintenance and preparation of L1 and L4 stages

The *C. elegans* wild type N2 strain was maintained at 20 °C on nematode growth medium (NGM) agar seeded with the *Escherichia coli* strain OP50. To prepare worm populations for screening, first a mixed population was obtained. 5 to 7 NGM plates were seeded with 250 μl of the *E. coli* strain OP50 and incubated at 37 °C for 3 to 5 days to form a lawn. A 2.5cm square section of the agar plate rich in worms was transferred to each NGM plate and maintained at 20 °C for 5 days.

Such *C. elegans* cultures contain worms of different ages. To eliminate variation caused by age differences, worms need to be synchronised. To obtain a synchronised L1 population we used filtration. Each NGM plate was washed with 50 mL of S-basal medium into a falcon tube and centrifuged at 3000 *g* at 20 °C for 4 min. The pellet was then re-washed (3x) using the same procedure to clear any remaining bacteria. Worms were then filtered (100 μm filter) to remove any adult and late-stage larvae. Finally, they were passed through a 40 μm sieve (3x) to obtain a synchronous L1 larval population.

To obtain a synchronised L4 larval population, a population enriched in L4s was first obtained by incubating the NGM plates, prepared as described, for between 8-10 days after transferring *C. elegans*. Plates with the highest number of L4 larvae were selected and washed with S-basal medium. Worms were then filtered (100 μm filter) to capture any L4 stage larvae and remove earlier stages. The filter was rinsed with S-basal media into a falcon tube and centrifuged at 3000 x *g* at 20 °C for 4 min. The pellet was then re-washed (2x) using the same procedure to remove any residual bacteria.

### The INVAPP / Paragon system for Automated Phenotyping of nematodes

The INVAPP / Paragon system was previously described (Partridge et al., 2018a; Buckingham et al., 2021). Briefly, the INVAPP / Paragon system consists of a high-resolution camera (Andor Neo, which captures images of 2560 × 2160 pixels) and a lens (Pentax YF3528). Microtiter (96 well) plates are placed in a holder built into the roof of an otherwise light-tight cabinet and imaged from below. Illumination is provided by an LED panel fitted with an acrylic diffuser.

Two hundred frame movies were captured at 25 frames s^-1^ for 8 s using μManager (Edelstein et al., 2014). Movies were analysed using MATLAB scripts (available at https://github.com/fpartridge/invapp-paragon) and the variance determined through time for each pixel. The distribution of these pixel variances was then considered, and pixels whose variance was above the threshold (those greater than one standard deviation away from the mean variance) were considered ‘motile’. Motile pixels within each well were counted to obtain a movement score. Full details of the device and its applications in worm motility and growth/motility screens have been published (Partridge et al., 2018a).

### *C. elegans* – growth/motility assay

Screening experiments were conducted in 96-well plate liquid cultures. Synchronised L1s were diluted to approximately 15-25 worms per 50 μl in S complete buffer with 1% w/v HB101 *E. coli*. Assay plates were prepared with 99 μL of L1 suspension and 1 μL of each drug compound per well. As a control 16 wells in each plate were prepared with 1 µL DMSO solution (1% v/v final concentration). Plates were incubated at 25 °C and motility was recorded using the INVAPP/Paragon system 3 days later. By this time control worms developed to L4 or adult stage. Movies were recorded and the median growth/motility score measured using INVAPP as described above. This assay measures both growth and motility in a combined score (Partridge et al., 2018a). A schematic of this assay is shown in Figure 1(A).

**Figure 1:**
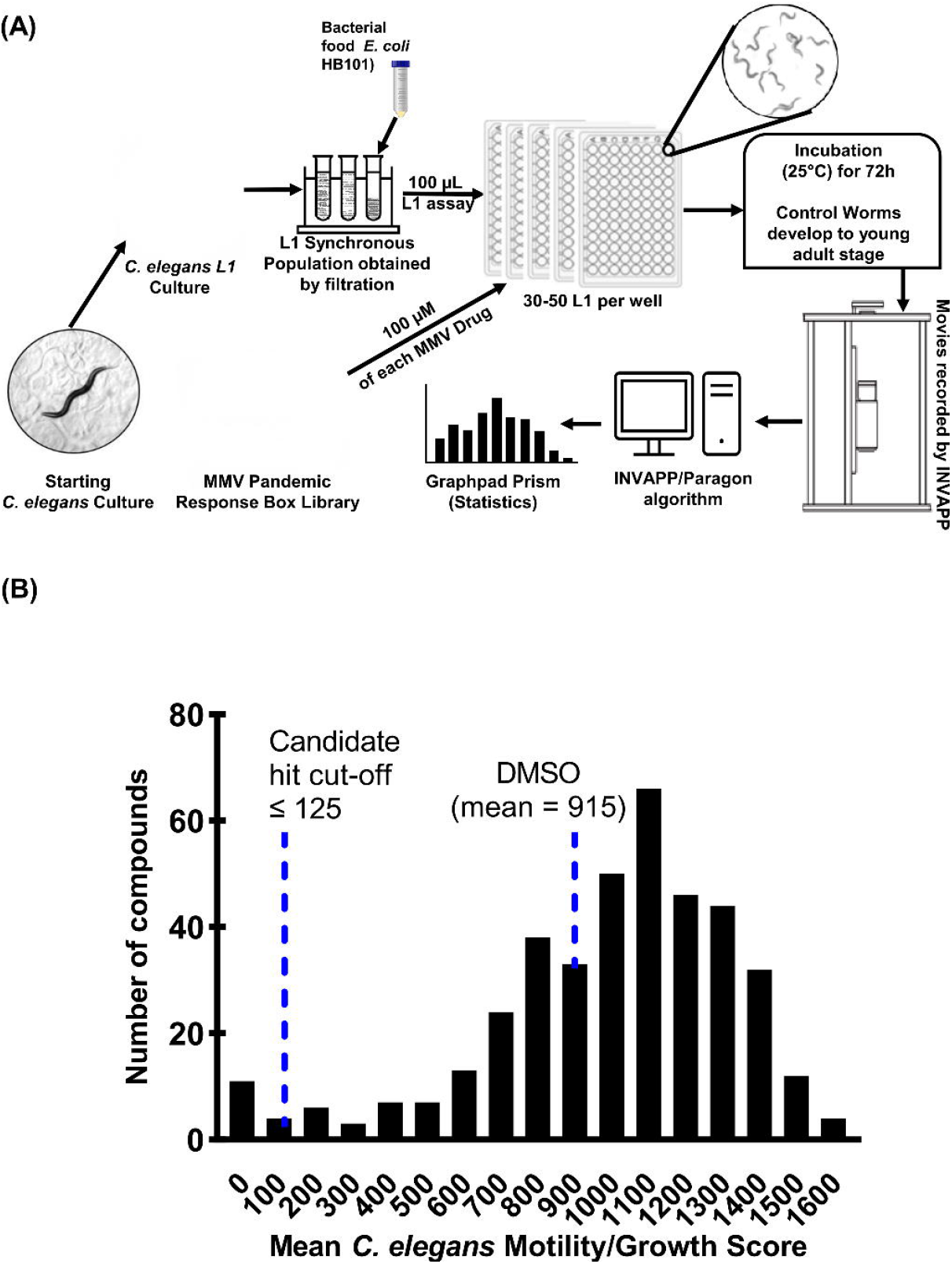
**(A)** Cartoon illustration of preparation and screening of *C. elegans* growth/motility assay. By the time that the movies are recorded, surviving *C. elegans* worms are expected to have developed to the (larger) L4 or adult stages. Therefore, their movement recorded by INVAPP/Paragon is a Growth/Motility score. **(B)** Frequency distribution of the mean *C. elegans* growth/motility scores for each of the Pandemic Response Box compounds tested at 1.0×10^−4^ M concentration, along with the DMSO-only control. Each bin is a range of scores from 50 below the axis value to 50 above the axis value. For example, the 200 bin counts the number of compounds scoring between 150 and 250. The screen was performed on three separate occasions (n=3). The mean of the three *C. elegans* growth/motility scores were plotted. A cut-off score for candidate hits of 125 was selected. The shaded region indicates compounds selected as candidate hits.

### *C. elegans* - motility assay

Synchronised L4 were diluted to approximately 15-25 worms per 50 μL in S basal buffer. No bacterial food is used in this assay. Assay plates were prepared with 99 μL of L4 suspension and 1 μL of each drug compound per well. As a control 16 wells in each plate were prepared with 1 µL DMSO solution (1% v/v final concentration). Plates were incubated at 25 °C before the motility of *C. elegans* was recorded using the INVAPP/Paragon system after 24 h. Due to the absence of food, worms do not grow or develop in this assay, which therefore exclusively measures changes in motility and is recorded as a motility score. A schematic of this assay is shown in Figure 3(A).

### Primary screen of MMV Pandemic Response Box library at 1.0×10^−4^ M using growth/motility assay

The 400 drugs of the MMV Pandemic Response Box library were screened at 1.0×10^−4^ M on wild type *C. elegans* in a growth/motility assay primary screen (L1 to L4 or adult development). 1% v/v DMSO was used as a negative control. Three sets of identical assay plates were prepared for each experiment and the entire screen was repeated on 3 different days (n=3).

### Secondary screen to confirm candidate lead compounds and additionally test activity in a motility assay

The 18 potential candidate lead compounds identified in the primary screen of the MMV Pandemic Response Box library were re-tested on *C. elegans* in the growth/motility assay, at 7.5×10^−5^ M (0.75% v/v DMSO) and 5.0×10^−5^ M (0.5% v/v DMSO). In addition, *C. elegans* L4 animals were screened at 1.0×10^−4^ M in the motility assay. In all cases, screens were undertaken on three separate occasions (n=3), each time with 4 assay repeats. Levamisole at the same concentration was the positive control and 1% v/v DMSO the negative control.

### Concentration-response curves

The concentration-response relationship for selected compounds was determined by testing activity in the same *C. elegans* growth/motility assay. Compounds were tested at each of 12 concentrations from 5.0×10^−5^ M to 2.0×10^−8^ M (10 to 12 replicates tested on three occasions so n=3). EC_50_ values were estimated by fitting curves using a four-parameter log-logistic function in Graphpad Prism 9.3.

## Results

### Identification of candidate lead compounds by screening the 400 compound MMV Pandemic Response Box Library in the *C. elegans* growth/motility assay

Primarily the Pandemic Response Box library was screened using the *C. elegans* growth/motility assay. The 400 compounds were screened at 1.0×10^−4^ M and the growth/motility score recorded (Partridge et al., 2018a). A histogram showing the distribution of the mean *C. elegans* growth/motility scores for each compound, along with the DMSO-only control is shown in Figure 1(B).

The primary screen was used to prioritise the most active compounds for confirmatory rescreening. We chose the 14 compounds with the lowest mean growth/motility score (3.5% of the library) for rescreening (each compound had a mean growth/motility score below 125). In addition, we selected a further 4 compounds for rescreening with scores close to this cut-off value. The full data for all three repeats of the primary screen using the *C. elegans* growth/motility assay at 1.0×10^−4^ M are presented in the Table S1.

### Confirmation of 18 active anthelmintic compounds in a secondary screen

To confirm the activity of the lead compounds, we conducted a secondary screen. The 18 candidates were re-tested in the same *C. elegans* growth/motility assay at two lower concentrations (7.5×10^−5^ M and 5.0×10^−5^ M). The results for the 7.5×10^−5^ M secondary screen are shown in Figure 2(A). A one-way ANOVA showed there was a significant effect of compound treatment (P ≤ 0.0001). The effectiveness of each compound compared to the DMSO-only control was determined using Dunnett’s multiple comparison test. All the 18 candidate lead compounds significantly reduced the *C. elegans* growth/ motility score at 7.5×10^−5^ M (Fig 2A, Table S3).

**Figure 2:**
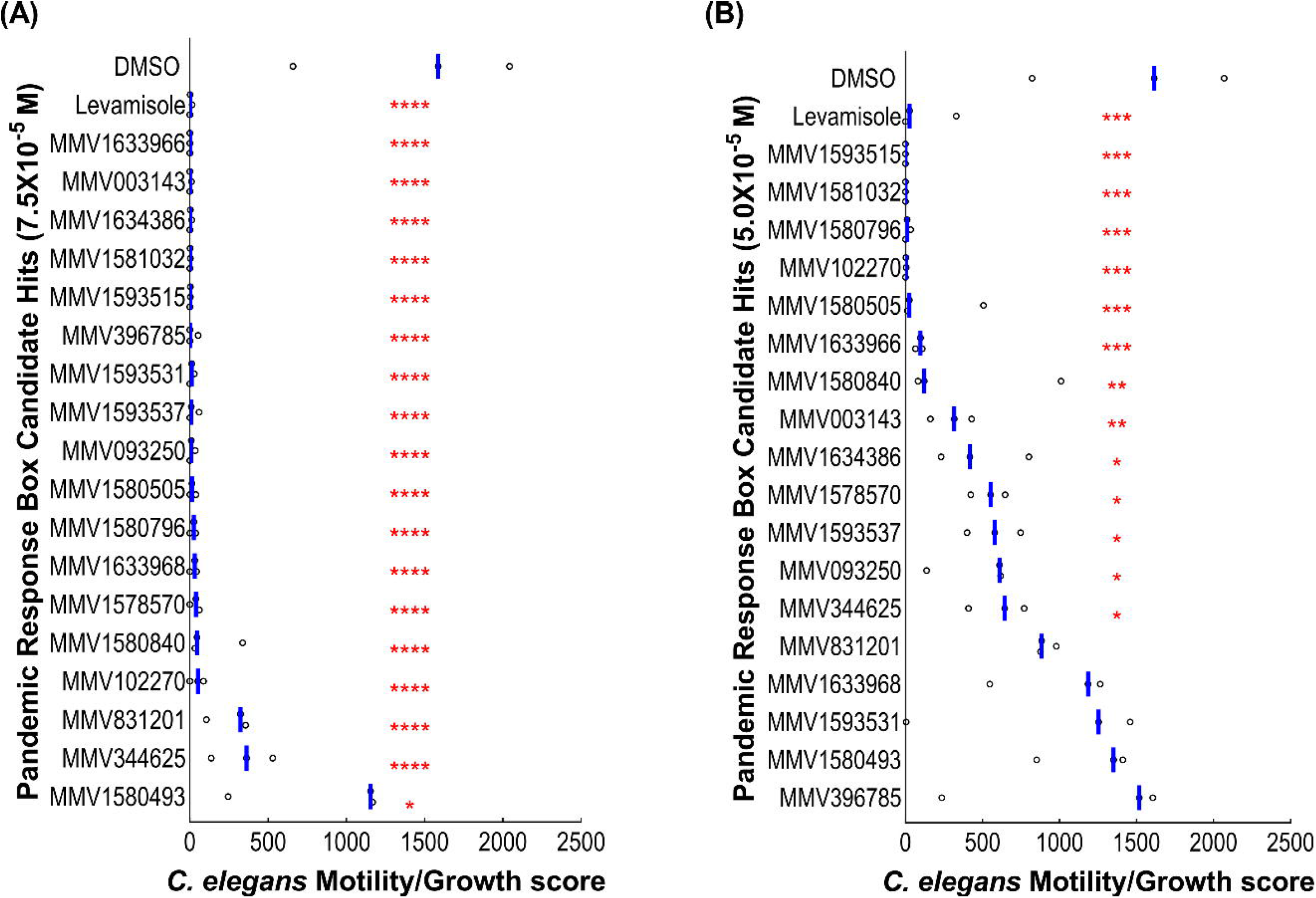
Retesting the 18 lead candidates as well as levamisole and DMSO as controls in the *C. elegans* growth/motility assay at **(A)** 7.5×10^−5^ M and **(B)** 5.0×10^−5^ M. Each point shows one well. The blue bar indicates the mean movement index for each treatment. The screen was performed with four replicates and on three occasions (n=3). A one-way ANOVA for the 7.5×10^−5^ M concentration dataset showed there was a significant effect of compound treatment P<0.0001. A post-hoc Dunnett’s test compared each compound to the DMSO-only control (P values* P ≤ 0.05, ** P ≤ 0.01, *** P ≤ 0.001, **** P ≤ 0.0001).

The results of the *C. elegans* growth/motility screen at 5.0×10^−5^ M are shown in figure 2(B). A one-way ANOVA showed when there was a significant effect of compound treatment (P ≤ 0.0001). The effectiveness of each compound was determined using Dunnett’s multiple comparison test with reference to the DMSO-only control. 13 Pandemic Response Box compounds significantly reduced the *C. elegans* growth/motility score at 5.0×10^−5^ M, as indicated by red asterisks in Figure 2(B). Six compounds were highly effective (P values ≤ 0.0001). Data for the secondary screen of the *C. elegans* growth/motility assay at 5.0×10^−5^ M are presented in Table S2.

### Identification of compounds that act to block motility

Many existing anthelmintics act by reducing motility, for example by acting on ion channels that function in the nervous system and /or at neuromuscular junctions (Raymond and Sattelle, 2002; Geary et al., 2015; Holden-Dye and Walker, 2018). We therefore wanted to determine which of the candidate lead compounds impair motility. The 18 candidate lead compounds were tested on *C. elegans* L4 stage worms in a pure motility assay at 1.0×10^−4^ M. This allowed us to verify whether, in addition to showing activity in the *C. elegans* growth/motility assay over 72 h, the candidate hit compounds were also effective in an assay using *C. elegans* L4 stage worms without bacterial food, over 24 h. The method is shown in Figure 3(A) and the results are presented in Figure 3(B). A one-way ANOVA for this dataset showed there was a significant effect of compound treatment (P ≤ 0.0001). The effectiveness of each compound was then determined using Dunnett’s multiple comparison test compared to the DMSO-only control. In addition to the positive control drug levamisole, 6 compounds that significantly reduced the *C. elegans* L4 motility score were identified and indicated with red asterisks (P values: * P ≤ 0.05, ** P ≤ 0.01). Data for the *C. elegans* L4 motility assay screen is presented in the Table S4.

**Figure 3:**
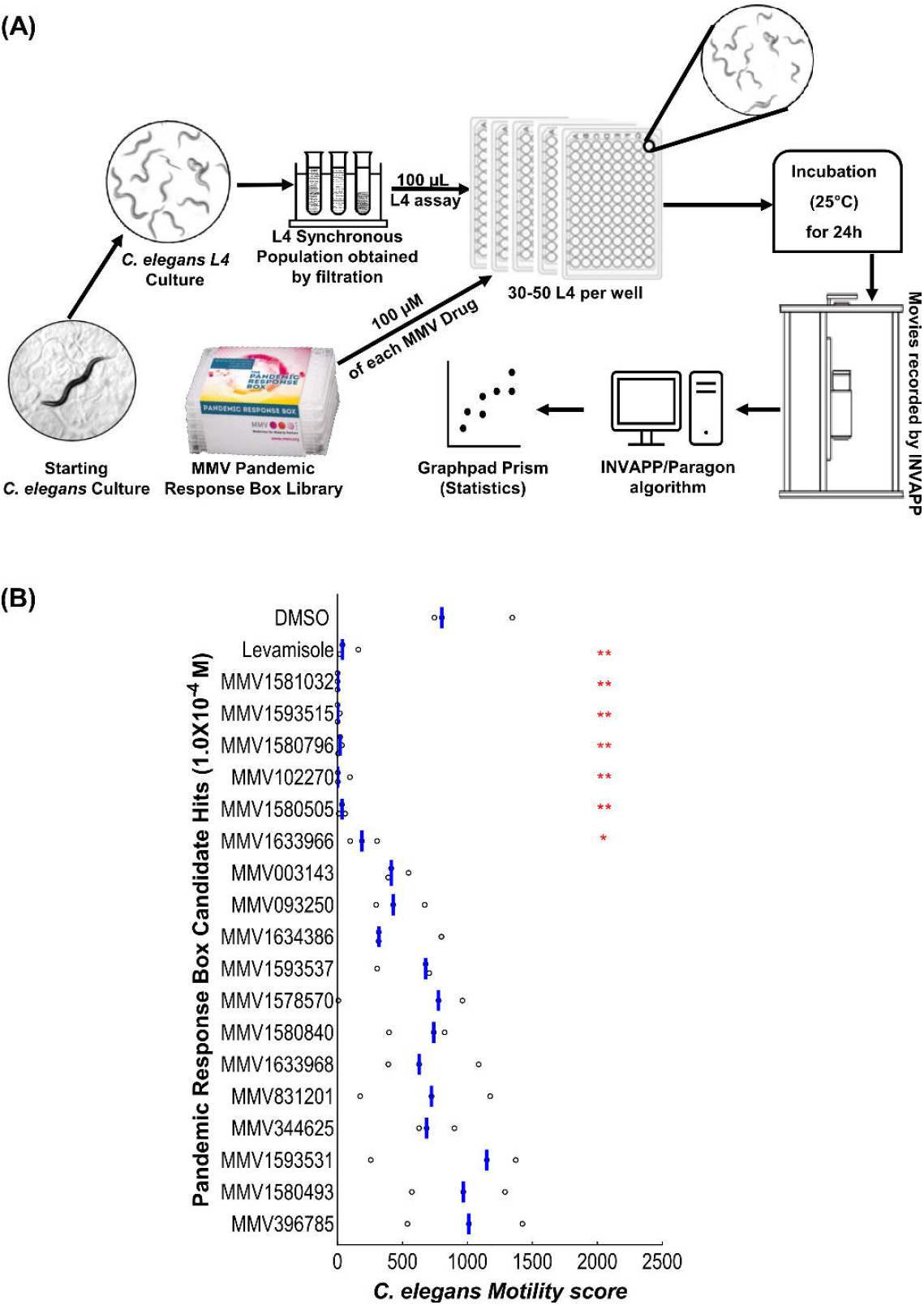
**(A)** Cartoon illustration of preparation and screening *C. elegans* motility assay. When the movies are recorded, surviving *C. elegans* worms have not developed further due to the lack of bacterial food. Therefore, their movement recorded by INVAPP/Paragon is a pure motility score. **(B)** Retesting the candidates in the *C. elegans* L4 motility assay at 1.0×10-4 M. Each point shows one well. The blue bar indicates the mean movement index for each treatment. The screen was performed with four replicates and on three occasions (n=3). A one-way ANOVA for the 1.0×10-4 M concentration dataset showed there was a significant effect of compound treatment (P ≤0.0001). A post-hoc Dunnett’s test compared each compound to the DMSO-only control (P values: * P ≤ 0.05, ** P ≤ 0.01).

### Summary of most active compounds in the secondary screens

Thus, six of the hit compounds from the Pandemic Response Box library were significantly active on both *C. elegans* growth/motility assay at 1. 0×10^−4^ M, 5.0×10^−5^ M and 7.5×10^−5^ M as well as the L4 motility assay at 1.0×10^−4^ M. These compounds were MMV1593515 (vorapaxar), MMV102270 (diphyllin), MMV1581032 (ABX464), MMV1580796 (rubitecan), MMV1580505 and MMV1593531. Four are previously described antivirals and two have antibacterial activity (Samby et al., 2022). Vorapaxar is a human approved drug while diphyllin, ABX464 and rubitecan are in various human clinical trials (Table 1).

**Table 1:**
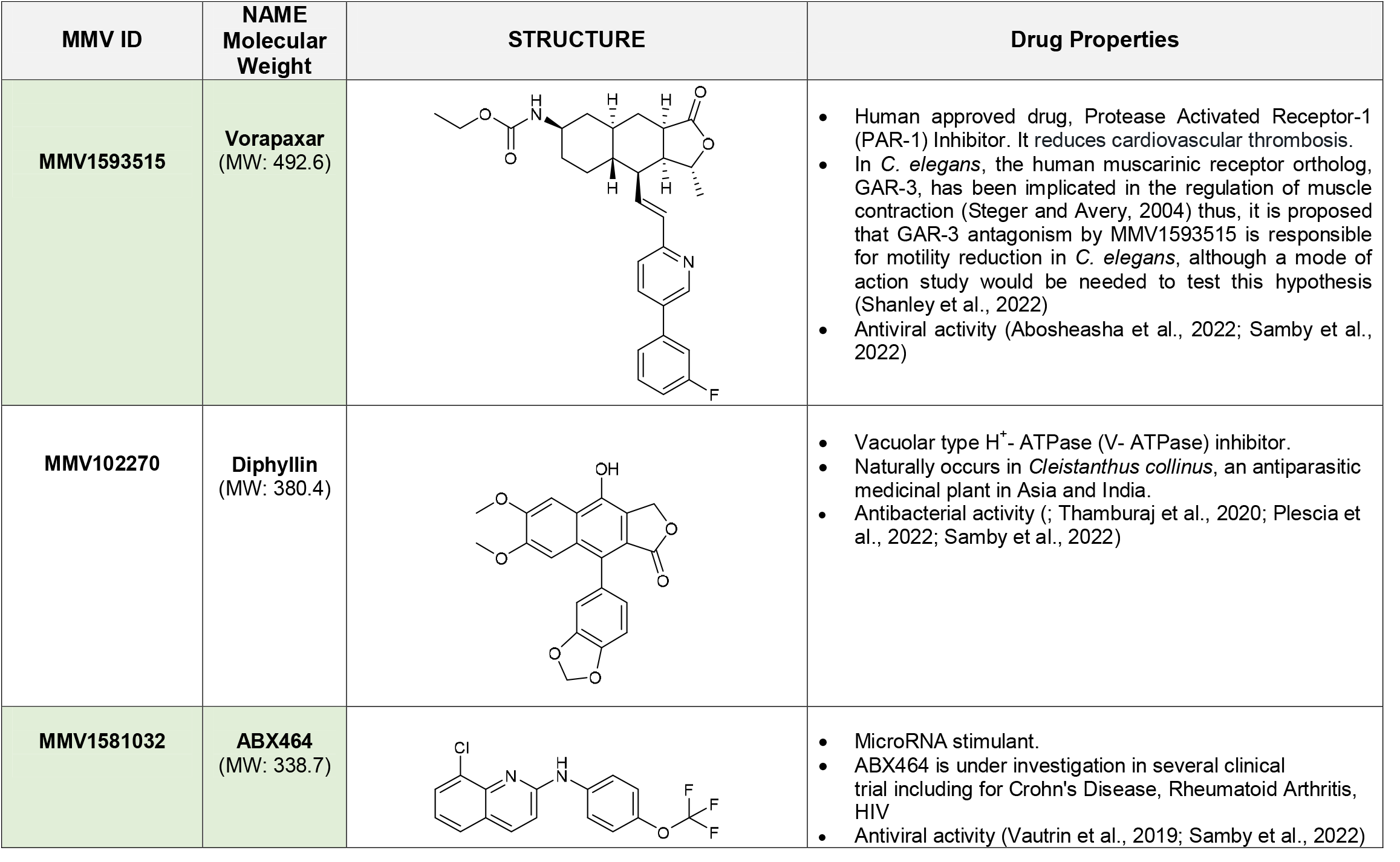
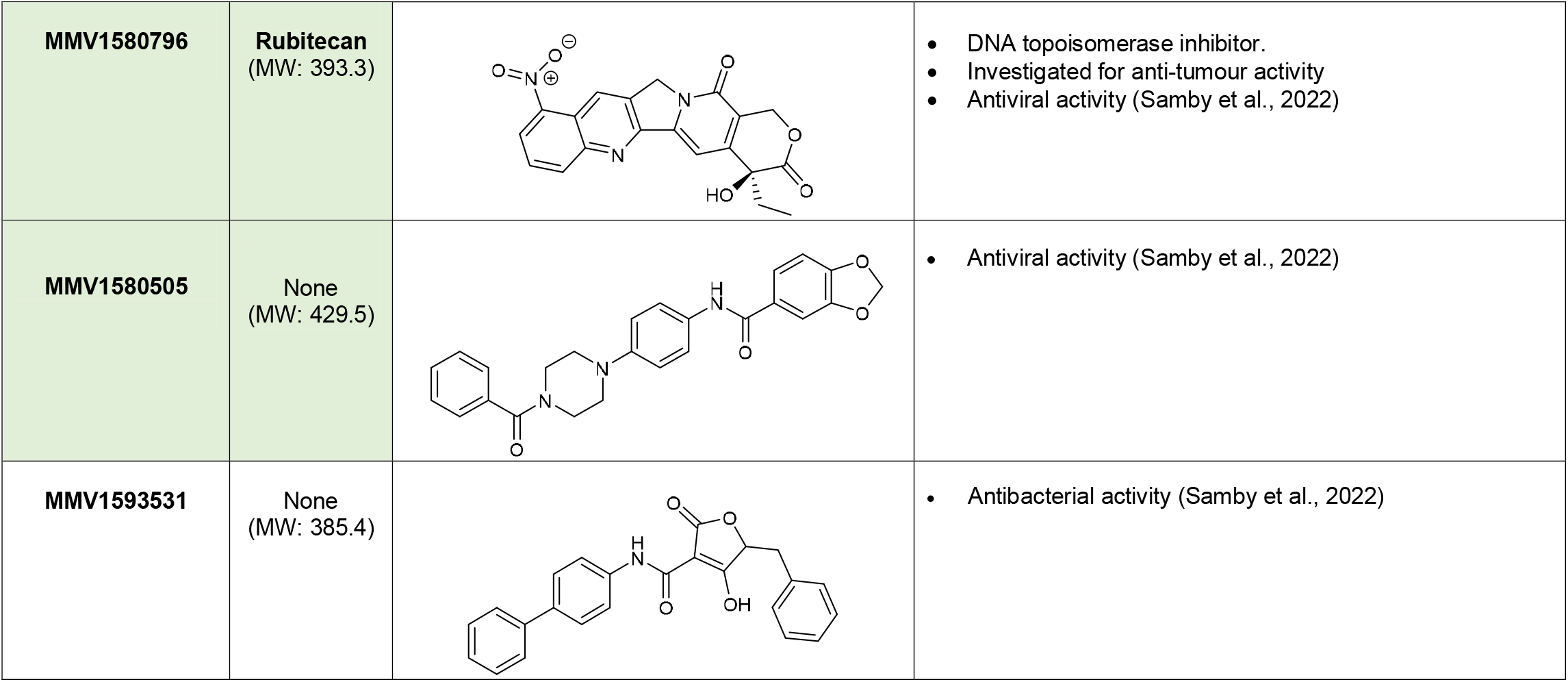
List of Pandemic Response Box hit compounds from the *C. elegans* growth/motility screen at 1.0×10^−4^ M, 5.0×10^−5^ M and 7.5×10^−5^ M and *C. elegans* L4 motility screen at 1.0×10^−4^ M. In this table the drugs highlighted with green shade are antiviral, and others are antibacterial drugs. Only vorapaxar is approved for human use.

| MMV ID | NAME<br>Molecular Weight | STRUCTURE | Drug Properties |
| --- | --- | --- | --- |
| MMV1593515 | Vorapaxar<br>(MW: 492.6) |  | Human approved drug, Protease Activated Receptor-1 (PAR-1) Inhibitor. It reduces cardiovascular thrombosis.<br>In *C. elegans*, the human muscarinic receptor ortholog, GAR-3, has been implicated in the regulation of muscle contraction (Steger and Avery, 2004) thus, it is proposed that GAR-3 antagonism by MMV1593515 is responsible for motility reduction in *C. elegans*, although a mode of action study would be needed to test this hypothesis (Shanley et al., 2022)<br>Antiviral activity (Abosheasha et al., 2022; Samby et al., 2022) |
| MMV102270 | Diphyllin<br>(MW: 380.4) |  | Vacuolar type H<sup>+</sup>-ATPase (V-ATPase) inhibitor.<br>Naturally occurs in *Cleistanthus collinus*, an antiparasitic medicinal plant in Asia and India.<br>Antibacterial activity (Thamburaj et al., 2020; Plescia et al., 2022; Samby et al., 2022) |
| MMV1581032 | ABX464<br>(MW: 338.7) |  | MicroRNA stimulant.<br>ABX464 is under investigation in several clinical trial including for Crohn's Disease, Rheumatoid Arthritis, HIV<br>Antiviral activity (Vautrin et al., 2019; Samby et al., 2022) |
| **MMV1580796** | **Rubitecan**<br>(MW: 393.3) |  | • DNA topoisomerase inhibitor.<br>• Investigated for anti-tumour activity<br>• Antiviral activity (Samby et al., 2022) |
| **MMV1580505** | None<br>(MW: 429.5) |  | • Antiviral activity (Samby et al., 2022) |
| **MMV1593531** | None<br>(MW: 385.4) |  | • Antibacterial activity (Samby et al., 2022) |

### Determination of the relative potency of the active anthelmintic compounds

We wanted to ensure that anthelmintic compounds we identified showed concentration-dependent activity, and to estimate their relative potency to inform future work. We used the efficacy of the best candidate lead compounds in the secondary screens to prioritize which to investigate. Of the six compounds significantly reducing L1 growth/motility at both 5.0×10^−5^ M and 7.5×10^−5^ M, as well as significantly reducing L4 motility, four (ABX464, diphyllin, rubitecan and vorapaxar) were readily available as solid material. Concentration-response curves for these four compounds are shown in Figure 4. It was encouraging to see EC_50_ values in the low micromolar – ABX464 (EC_50_ = 2.3 × 10^−6^ M), diphyllin (EC_50_ = 3.9 × 10^−6^ M) and rubitecan (EC_50_ = 1.2 ×10^−5^ M) and even sub-micromolar range – vorapaxar (EC_50_ = 5.7 × 10^−7^ M).

**Figure 4:**
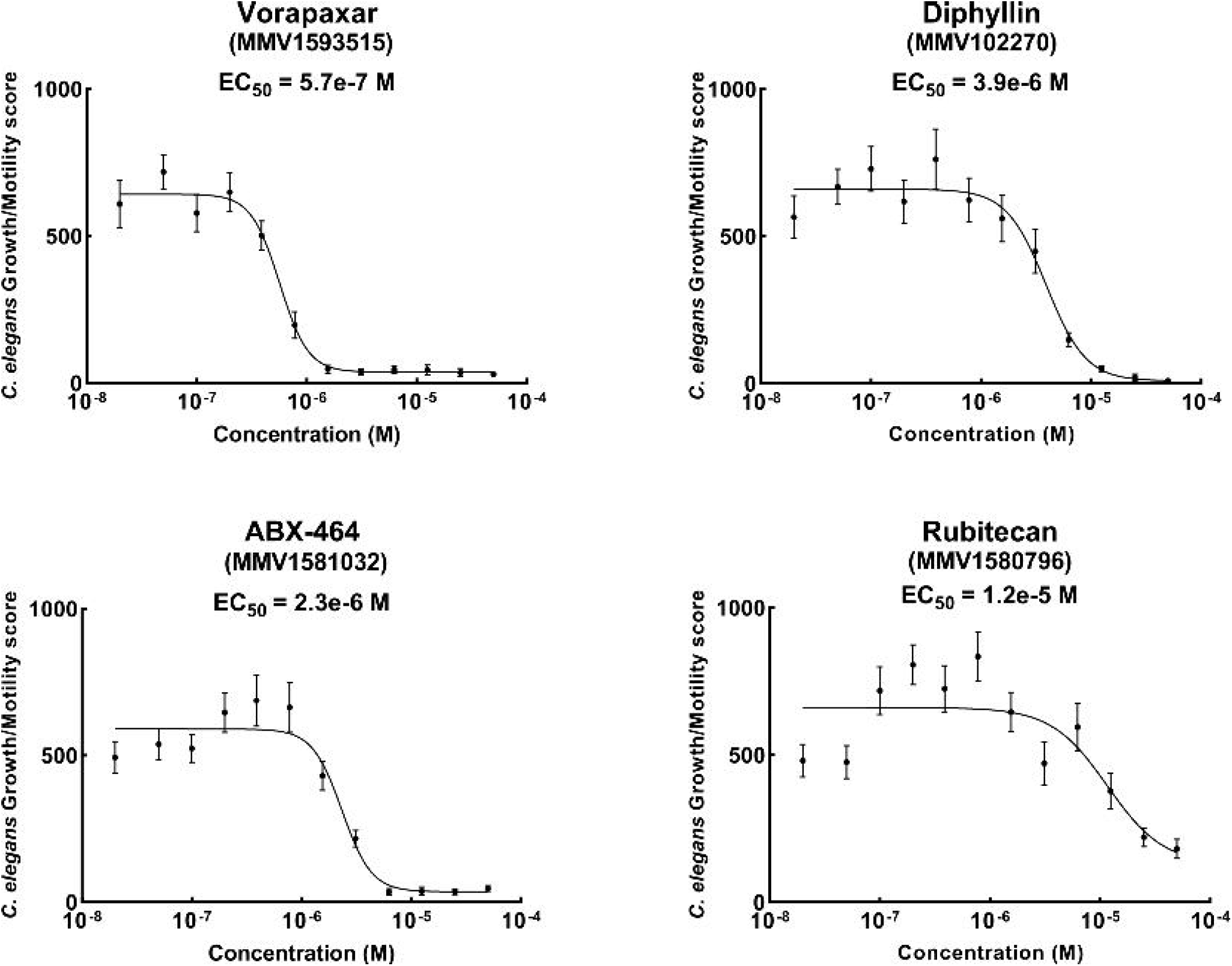
EC_50_ values are determined from concentration-response curves for the 4 compounds consistently showing activity in the growth/motility assay. Each point shows the mean and standard error (error bars) for each concentration tested. Compounds were tested between 10 to 12 replicates and 12 concentrations ranging from 5.0×10^−5^ M to 2.0×10^−8^ M (tested on three occasions so n=3). EC_50_ curves were fitted using a four-parameter log-logistic function in Graphpad Prism 9.3.

In mammalian studies, MMV1593515 (vorapaxar) the human approved drug is a protease activated receptor-1 (PAR-1) inhibitor (Heuberger and Schuepbach, 2019; Zhang et al., 2020), MMV102270 (diphyllin) is a vacuolar type H^+^-ATPase (V-ATPase) inhibitor (Holliday, 2017; Chen et al., 2018), MMV1581032 (ABX464) is a microRNA stimulant (Tazi et al., 2021) and MMV1580796 (rubitecan) is a DNA topoisomerase inhibitor (Burris et al., 2005). Thus the 4 candidate compounds of interest have diverse mechanisms of action but all showed anthelmintic activity. MMV1593515 (vorapaxar), MMV1581032 (ABX464) and MMV1580796 (rubitecan) are antivirals and MMV102270 (diphyllin) is an antibacterial drug. Their mechanisms of action in nematodes remain to be determined.

## Discussion

The 400 compound MMV Pandemic Response Box library has proved useful in identifying from library screens compounds of interest for re-purposing as antiparasitic for helminth control, and the control of schistosome and protozoan parasites. A recent study reported a screen of the MMV Pandemic Response Box against *C. elegans* young adults in a motility screening assay. This assay is similar to our L4 motility assay that is reported in Figure 3. They also screened against *Haemonchus contortus* exsheathed third-stage larvae (xL3s) in motility and development screening assays (Shanley et al., 2022).

Shanley *et al* identified two compounds, MMV1581032 (ABX464) and MMV1593515 (vorapaxar), that inhibited *C. elegans* motility. These compounds are among our four most active compounds, and we confirm their relatively low EC_50_ values.

In addition, Shanley et al (2022) showed that MMV1581032 (ABX464), as well as another drug, MMV1593539, have anthelmintic activity against *H. contortus*. As in their studies, we also found that MMV1593539 is not active against *C. elegans*. The accord between our findings and those of Shanley et al (2022) confirm the utility of our INVAPP screening approach.

However, in this study we also identify an additional 16 active anthelmintic compounds. This difference is likely due to our use of a *C. elegans* assay that measures growth/motility as worms develop from the L1 stage. This might reflect either a greater sensitivity of the L1 stages, or the choice of a higher initial screening concentration. Differential sensitivity of different life stages of nematodes and the importance of this in the choice of screening assays in order to avoid false negatives has been well described elsewhere (Elfawal et al., 2019). There are likely more biological processes (and hence drug targets) that could be targeted by anthelmintic assays in our growth/motility assay compared to a pure motility assay, which may account for the greater diversity of compounds detected with anthelmintic properties. The known inactivity of the approved anthelmintic drug mebendazole on L4 motility in *C. elegans*, compared to its high activity in the L1 growth/motility assay (Partridge et al., 2018a) demonstrates that viable anthelmintics will be missed by pure motility screens. Our study demonstrates the utility of INVAPP / Paragon and is complementary to that of Shanley *et al*. (2022).

MMV1581032 (ABX464) is a first-in-class, clinical-stage, oral small molecule immunomodulator (Vautrin et al., 2019). ABX464 is reported to bind to the RNA cap-binding complex, which modulates both viral and cellular RNA biogenesis (Tazi et al., 2021). ABX464 was originally developed for its antiviral potential but it was redeployed for chronic inflammatory diseases due to potent anti-inflammatory effects in preclinical testing. It has shown safety and tolerability in clinical trials, which makes it a good candidate for repurposing as an anthelmintic.

Diphyllin occurs naturally in *Cleistanthus collinus*, which is known as an antiparasitic medicinal plant in Asia and India (Suman et al., 2018). Diphyllin proved deadly for the promastigote and amastigote stages of the protozoan parasite *Leishmania infantum* (Di Giorgio et al., 2005), while the structurally related compound justicidin β suppressed the growth of the parasites causing sleeping sickness *Trypanosoma brucei rhodesiense* and *T. cruzi* (Gertsch et al., 2003).

The MMV Pandemic Response Box library has also been screened against *Schistosoma mansoni* with the aim of discovering an effective drug for the neglected tropical disease schistosomiasis (Biendl et al., 2022). The study identified 26 compounds active against newly transformed schistosomula, of which 17 were active against adult *S. mansoni*. Three compounds with anti-schistosomal activity (MMV396785 [Alexidine], MMV1634386 and MMV1578570) showed activity against *C. elegans* growth/motility in this study. These compounds therefore have potential for development as a broad-spectrum anthelmintics.

Some of our confirmed active compounds have also shown activity against other infectious diseases. MMV1593537 has antifungal activity against *Cryptococcus neoformans, Cryptococcus deuterogattii*, and the emerging global threat *Candida auris* (de Oliveira et al., 2022). MMV1578570, MMV396785 (alexidine), MMV1634386 and MMV1580796 (rubitecan) are active against the pathogenic amoebae *Balamuthia mandrillaris, Naegleria fowleri, and Acanthamoeba castellanii* (Rice et al., 2020). Rubitecan is one of our four most active anthelmintic compounds. It prevents DNA from unwinding during replication via DNA topoisomerase 1, therefore interfering with tumour growth (Lian et al., 2014). Rubitecan is a derivative of a compound extracted from the *Camptotheca acuminata* tree with potent antitumor and antiviral properties.

## Conclusion

This study identified 18 candidate lead compounds which impair nematode growth or motility, and the four most active include one drug (vorapaxar) with a sub-micromolar EC_50_, comparable to some current commercial anthelmintic drugs. The modes of action of the Pandemic Response Box hit compounds in nematodes are currently unknown, although it might be investigated utilising genomic, transcriptomic, and proteomic approaches like those used to uncover the molecular target of the veterinary anthelmintic monepantel.

## Acknowledgements

The authors would like to thank Medicines for Malaria Venture (MMV) for supplying the Pandemic Response Box.

## Supplementary Material

The following files are available online

**S1 Table:**

Result of **Primary screen** of MMV Pandemic Response Box library at **1.0×10**^**-4**^ **M** using growth/motility assay and DMSO-only as negative control. Table S1 contains data for three separate screen batches undertaken at three separate occasions each with 3 x identical replicates for each compound. INVAPP score indicates the rate of movement for each well of 96 well plates containing growth/motility assay. The table also contains the position of each compound at the experiments. Median rate for each replicate also included in the table with MMV name of each drug, Trival name, Disease area and Smile structure of each compound. The cut off to identify active compounds also presented.

**S2 Table:**

Result of **Secondary screen** of the 18 hit compounds identified in the primary screen of the MMV Pandemic Response Box library. They were re-tested on *C. elegans* L1s growth/motility assay at **5.0×10**^**-5**^ **M** and (0.5% v/v DMSO) as negative control. Screens were undertaken at 3 separate occasions with 4 replicates each (n=3) with similar concentration of levamisole as a positive control. INVAPP score indicates the rate of movement for each well of 96 well plates containing growth/motility assay. Table also contains the position of each compound at the experiments, median rate for each replicate with MMV name of each compound.

**S3 Table:**

Result of **Secondary screen** of the 18 hit compounds identified in the primary screen of the MMV Pandemic Response Box library. They were re-tested on *C. elegans* L1s growth/motility assay at **7.5×10**^**-5**^ **M** and (0.75% v/v DMSO) as negative control. Screens were undertaken at 3 separate occasions with 4 replicates each (n=3) with similar concentration of levamisole as a positive control. INVAPP score indicates the rate of movement for each well of 96 well plates containing growth/motility assay. Table also contains the position of each compound at the experiments, median rate for each replicate with MMV name of each compound.

**S4 Table:**

Result of **Secondary screen** of the 18 hit compounds identified in the primary screen of the MMV Pandemic Response Box library. They were re-tested on *C. elegans* L4 motility assay at **1.0×10**^**-4**^ **M** and DMSO-only as negative control. Screens were undertaken at 3 separate occasions with 4 replicates each (n=3) with similar concentration of levamisole as a positive control. INVAPP score indicates the rate of movement for each well of 96 well plates containing motility assay. Table also contains the position of each compound at the experiments, median rate for each replicate with MMV name of each compound.

